# Analysis of the off-target interaction of amyloid PET tracers with human brain sulfotransferases

**DOI:** 10.64898/2026.06.16.732640

**Authors:** Ludovico Miccoli, Rosa Fullone, Stefano Delli Pizzi, Federica Tomaiuolo, Stefano L. Sensi, Giuseppe Floresta, Alberto Granzotto

## Abstract

Positron emission tomography (PET) tracers targeting amyloid-β (Aβ) are central to the diagnosis and staging of Alzheimer’s disease (AD). However, growing evidence indicates that these tracers can engage off-target molecules, complicating signal interpretation. Sulfotransferases (SULTs) have been experimentally identified as binding partners of ^11^C-Pittsburgh Compound-B (PiB). However, whether the clinically used fluorinated PiB derivatives flutemetamol and flutafuranol interact with brain-expressed SULTs is yet unexplored. Here, we combined multi-omic transcriptomic profiling with molecular docking and molecular dynamics (MD) simulations to assess the structural interactions of SULT-tracer complexes. Analysis of the Genotype-Tissue Expression project and the Human Protein Atlas identified SULT1A1, SULT1A3, and SULT4A1 as the SULT isoforms predominantly expressed in the human brain. Docking and MD simulations showed that all three tracers form energetically comparable complexes within the catalytic pockets of these isoforms, yet their dynamic stability varied in an enzyme-and tracer-specific manner. PiB and flutemetamol were stably accommodated in SULT1A1, but PiB lost its initial pose in SULT4A1. Flutafuranol showed weaker binding to SULT1A1, yet formed stable complexes in SULT1A3 and SULT4A1. Notably, SULT1A1, SULT1A3, and SULT4A1 are all expressed in the cerebellum, the brain region used as a reference for Aβ PET signal normalization. These findings provide a structural framework for off-target tracer interaction with brain SULTs and suggest that the intracellular enzymatic environment may contribute to variability in Aβ PET signals beyond fibrillar Aβ deposition.

## Introduction

Alzheimer’s disease (AD) is one of the leading causes of dementia, affecting over 55 million people worldwide [1]. This figure is expected to nearly triple by 2050, imposing an escalating socioeconomic burden [2]. Early identification of individuals at risk of developing AD is therefore imperative for the timely implementation of disease-modifying interventions [3]. The recent shift from a clinical to a biological definition of AD, as formalized by the ATN framework (i.e.,’A’ – Aβ,’T’ – tau, and’N’ – neurodegeneration), underscores the central role of biomarkers in diagnosis and disease monitoring [4]. In this context, Aβ positron emission tomography (PET) tracers provide a unique opportunity to directly visualize pathological β-amyloid (Aβ) deposition in the living human brain, one of the earliest events in the AD continuum [4, 5].

^11^C-Pittsburgh Compound-B (PiB) was the first tracer successfully employed to detect Aβ deposits in humans [6]. PiB is based on the benzothiazole backbone of thioflavin-T (ThT) [7]. However, its short radioactive half-life led to the development of fluorine-18 (^18^F) derivatives, such as flutemetamol [8] and flutafuranol (NAV4694/AZD4694) [9]. The latter possesses a distinct benzofuran-based structure offering higher affinity for Aβ deposits and reduced off-target interactions and non-specific white matter binding [9]. While the use of PiB is limited to specialized research settings, flutemetamol and flutafuranol - along with the trans-stilbene-based compounds ^18^F-florbetapir and ^18^F-florbetaben - are widely employed for the diagnosis of AD and for monitoring the efficacy of Aβ-targeting interventions [5]. Despite their widespread adoption and integration into diagnostic criteria and disease staging [4, 5], the specificity of these tracers remains a subject of debate [5, 9–12]. Although PiB and its derivatives were designed to bind fibrillar Aβ with high affinity, non-specific binding has been reported in white matter [13], meninges [14, 15], vascular structures [16], and the skull [17, 18]. Furthermore, more recently, atypical off-target signals have been reported in Aβ PET scans using NAV4694, including tracer uptake in the venous sinuses and choroid plexus, which can interfere with quantitative and visual interpretation [19]. Collectively, these observations reinforce the need to account for potential off-target binding when interpreting Aβ PET images.

Sulfotransferases (SULTs) are a class of enzymes that catalyze the sulfonation of neurotransmitters, hormones, and xenobiotics by transferring a sulfonate group from the donor 3′-phosphoadenosine-5′-phosphosulfate (PAPS) to various endogenous and exogenous substrates [20]. SULT4A1 is a brain-enriched SULT isoform whose activity has been linked to neurodevelopment and to the metabolism of neurotransmitters such as dopamine [21]. Recently, SULT4A1 has been also implicated in synaptic stability and in the modulation of the PSD-95/NMDAR complex [22]. Expression of other SULT isoforms, like SULT1A1, SULT1A3, and SULT1E1, has also been reported in the central nervous system (CNS) [23, 24]. *In vitro* and *in vivo* experimental work has established SULT1E1 as a *bona fide* PiB-binding partner [23, 25], demonstrating that 2-aryl-6-hydroxybenzothiazole-based tracers can stably interact with the SULT catalytic pocket. Whether this binding mode extends to SULT isoforms abundantly expressed in the human brain – and whether the clinically used fluorinated PiB derivatives share this off-target liability – remains unexplored.

This study evaluates the stability and topological relevance of interactions between brain SULTs and PiB as well as its fluorinated derivatives. We first assessed the brain expression of SULT isoforms through a systematic analysis of publicly available transcriptomic datasets, focusing on SULTs found most consistently detected in the human brain. Next, we integrated the expression profiles with structure-based *in silico* analysis by combining molecular docking with molecular dynamics (MD) simulations. This integrated approach is instrumental to evaluate whether specific SULTs could constitute a yet underappreciated source of off-target binding in Aβ PET imaging of AD.

## Materials and Methods

### Identification of brain enriched SULT isoforms

To establish a biological rationale for the potential interaction between Aβ-PET tracers and human SULTs, we performed a systematic multi-omic characterization of SULT1A1, SULT1A3, SULT1E1, and SULT4A1. Quantitative mRNA expression data (expressed as Transcripts Per Million, TPM) were retrieved from the Genotype-Tissue Expression (GTEx) project (v8), leveraging RNA-sequencing (RNA-seq) data from 54 tissue sites and 13 brain subregions [26]. To further support transcript expression at the cellular level, we integrated transcriptomic evidence from the Human Protein Atlas (HPA) [27]. This approach enabled the assessment of SULT transcript stability and localization.

### Regional expression data

To characterize the regional cortical distribution of SULT isoforms, we used transcriptomic data from the cortical transcriptional cartography dataset described in [28]. This resource integrates post-mortem gene expression profiles from six donors of the Allen Human Brain Atlas and projects them onto a standardized cortical surface in fsLR 32k space, providing vertex-wise maps of transcriptional variation across the human cortex. Briefly, cortical expression values represent within-subject z-scored gene expression averaged across donors, thereby reflecting the relative spatial distribution of each transcript rather than absolute quantitative differences across genes. For each SULT isoform, vertex-wise expression data were extracted, converted into GIFTI surface files, and resampled from fsLR to fsaverage space for visualization. Maps were displayed on the fsaverage pial surface while preserving the original spatial topology. No additional spatial smoothing, thresholding, or inter-gene normalization was applied.

### Ligand and Protein Preparation

The chemical structures of PiB, flutemetamol, and flutafuranol were retrieved from the PubChem database as three-dimensional conformers in Structure Data File (.sdf) format (^18^F-Flutemetamol, PubChem CID: 15950376; 11C-PiB, CID: 2826731; 18F-Flutafuranol, CID: 44815683).

Crystal structures of human brain-expressed SULT isoforms were obtained from the Protein Data Bank (PDB), including SULT1A1 (PDB ID: 2D06), SULT1E1 (PDB ID: 4JVL), and SULT1A3 (PDB ID: 2A3R). All protein structures were subjected to restrained energy minimization using the AMBER14 force field as implemented in YASARA to relieve steric clashes, optimize side-chain orientations, and improve local geometry while preserving the overall backbone conformation.

For SULT4A1, for which no high-resolution crystallographic structure is currently available, the predicted structure was obtained from the AlphaFold Protein Structure Database (entry AF-Q9BR01-F1-v4), characterized by a high average predicted Local Distance Difference Test (pLDDT) score of 87.22, indicating reliable backbone and side-chain positioning.

### Molecular Docking and Interaction Analysis

Molecular docking simulations were performed using the AutoDock Vina algorithm integrated within the YASARA Structure suite. Prior to docking, co-crystallized ligands, ions, and non-structural water molecules were removed from the protein structures. The docking search space was defined as a three-dimensional grid encompassing the entire active site cavity and surrounding residues potentially involved in ligand recognition, ensuring complete exploration of the binding region. Docking calculations were carried out using the default Vina scoring function and search parameters, followed by local refinement using the YASARA implementation of the Vina local search protocol to improve ligand placement and interaction geometry. For each ligand–protein pair, multiple binding poses were generated and ranked according to predicted binding affinity (ΔGbind, kcal/mol). Two alternative binding orientations were evaluated for each ligand, defined by the position of the benzo-fused heterocycle relative to the catalytic site. A standardized alphanumeric nomenclature was adopted to identify tracer identity and binding orientation. Tracers were assigned numerical identifiers: PiB (1), flutemetamol (2), and flutafuranol (3). Binding orientation was indicated by alphabetical suffixes, where “a” denotes the catalytic site–oriented configuration, in which the benzo-fused heterocycle is positioned proximal to the catalytic site, and “b” denotes the solvent-oriented configuration, in which the benzo-fused heterocycle is oriented toward the solvent-exposed region of the binding pocket. Both orientations were subsequently subjected to MD simulations to evaluate their structural stability.

The protein-ligand interaction networks were analyzed and visualized using BIOVIA Discovery Studio Modeling Environment [29]. This platform was employed to generate two-dimensional (2D) interaction diagrams, allowing for the precise characterization of non-covalent binding motifs, including conventional hydrogen bonds, π-stacking interactions, sulfur-π contacts, and hydrophobic networks. These analyses were critical for identifying the most favorable docking poses and for monitoring the maintenance of key stabilizing interactions throughout the subsequent MD trajectories.

### MD Simulations

All selected protein–ligand complexes were subjected to MD simulations using the YASARA Structure package and the md_runfast.mcr macro. Each complex was placed in a periodic cuboid simulation box extending at least 10 Å from any atom of the solute, ensuring adequate solvation and avoiding boundary artifacts. The simulation cell was filled with explicit TIP3P water molecules at a density of 0.997 g/mL. Physiological ionic strength was reproduced by adding Na⁺ and Cl⁻ ions at a final concentration of 0.9% (w/v), and overall charge neutrality was ensured. System preparation included optimization of the hydrogen-bonding network and prediction of residue pKa values to assign appropriate protonation states at physiological pH (7.4). Production MD simulations were performed for 300 ns using the AMBER14 force field for protein atoms, GAFF2 with AM1BCC charges for ligand parametrization, and TIP3P for explicit solvent molecules. Long-range electrostatic interactions were treated using the Particle Mesh Ewald (PME) algorithm without cutoff, while van der Waals interactions were calculated using an 8 Å cutoff radius with appropriate smoothing functions. The equations of motion were integrated using a multiple time-step algorithm, employing a time step of 2.5 fs for bonded interactions and 5.0 fs for non-bonded interactions. Simulations were conducted under constant temperature and pressure conditions (298 K and 1 atm) using the NPT ensemble, with temperature and pressure regulated using Berendsen-type coupling algorithms as implemented in YASARA. Trajectory snapshots were recorded every 250 ps for subsequent structural and energetic analyses.

### Trajectory Analysis and Binding Energy Calculations

Trajectory analysis was performed using built-in YASARA analysis macros. Structural stability of the protein–ligand complexes was evaluated by calculating the root-mean-square deviation (RMSD) of ligand heavy atoms after least-squares fitting of the protein backbone to the initial structure, allowing assessment of ligand positional stability within the binding pocket over time. Binding free energies were calculated using the Molecular Mechanics/Poisson–Boltzmann Surface Area (MM/PBSA) method, implemented in the md_analyzebindenergy macro of YASARA. This approach estimates the binding free energy by combining molecular mechanics interaction energies, polar solvation energies obtained from Poisson–Boltzmann calculations, and nonpolar solvation contributions based on solvent-accessible surface area. Representative structures, interaction patterns, and binding energy trajectories were extracted from equilibrated portions of the simulations for comparative analysis across the different SULT isoforms and ligands. To ensure equilibration, the first 50 ns of each trajectory were excluded, and analyses were performed on the remaining 250 ns. Statistical parameters derived from RMSD calculations are reported in Sup. Fig. 2.

## Results

### Brain expression and spatial mapping of SULT isoforms

SULTs include at least thirteen distinct members across three major families, SULT1, SULT2, and SULT4 [20]. To identify the isoforms primarily expressed in the human brain, we interrogated two independent transcriptomic databases: the Genotype-Tissue Expression (GTEx) project database [26] and the Human Protein Atlas [27]. Qualitative assessment of SULT expression levels showed that SULT4A1 is the primary isoform expressed in the human cortex (Sup. Fig. 1). SULT1A1 and SULT1A3 were also abundantly expressed (Sup. Fig. 1). Contrary to previous reports, expression levels of SULT1E1 were negligible (Sup. Fig. 1). Transcriptomic data from the Human Protein Atlas confirmed the presence of SULT1A1, SULT1A3, and SULT4A1 in the human cortex. Notably, the three isoforms were also abundantly expressed in the cerebellum.

To further characterize the regional cortical distribution of SULT isoforms, we examined surface-based transcriptional brain maps derived from the cortical transcriptional cartography dataset described in [29]. Expression patterns are displayed as vertex-wise within-gene z-scored maps (Sup. Fig. 1C), reflecting the relative spatial distribution of each transcript rather than absolute expression differences across genes. Relative cortical enrichment of SULT4A1 transcripts was observed in parietal and occipital regions, whereas lower relative representation was evident in ventral temporal areas and the anterior frontal cortex. SULT1A3 showed relative enrichment predominantly in occipital regions, with intermediate representation in the middle temporal gyrus and lower levels across frontal areas. SULT1A1 exhibited its highest relative cortical representation within the parietal cortex, with additional enrichment in anterior temporal regions, while occipital and frontal cortices showed comparatively lower representation.

Taken together, transcriptomic profiling and cortical mapping analyses indicate that SULT4A1, SULT1A1, and SULT1A3 are the predominant SULT isoforms expressed in the human brain, each displaying a distinct spatial cortical distribution. In contrast to previous reports, SULT1E1 expression was negligible across the datasets analyzed.

### Docking-derived binding orientations and MD simulation analysis

To evaluate the dynamic behavior of tracer-protein interactions, we performed local docking of PiB (1), flutemetamol (2), and flutafuranol (3) within the catalytic pockets of SULT1A1, SULT1A3, and SULT4A1. SULT1E1 was included as a reference isoform given that its interaction with PiB has been previously validated through *in vitro* binding assays and *in vivo* ^11^C-PiB autoradiography [23, 25], providing an experimentally validated benchmark. Docking analysis identified two energetically comparable ligand orientations for SULT1A1, SULT1E1, and SULT1A3. The two orientations primarily differed in whether the benzo-fused heterocycle was oriented inward (a) or outward (b) relative to the catalytic site. In contrast, docking to SULT4A1 consistently yielded a single pose in which the benzo-fused heterocycle was oriented toward the catalytic site.

Each pose was subsequently subjected to a 300 ns MD simulation. Dynamic stability was quantified using three complementary metrics: (i) the mean RMSD, (ii) the RMSD standard deviation (SD), and (iii) the RMSD slope over time. Configurations with near-zero RMSD slopes were prioritized as suggestive of stable binding modes [30, 31]. The prioritized poses were then validated by analysis of ligand–residue interactions within the catalytic pocket. With this rationale in mind, the final orientations analyzed were: SULT1E1-PiB (1a), SULT1E1-flutemetamol (2b), SULT1E1-flutafuranol (3b), SULT1A1-PiB (1b), SULT1A1-flutemetamol (2a), SULT1A1-flutafuranol (3a), SULT1A3-PiB (1a), SULT1A3-flutemetamol (2a), and SULT1A3-flutafuranol (3a) (Sup. Fig. 2; Sup. Tab. 1). Within the SULT1E1 binding pocket, PiB (1a) forms hydrogen bonds with His A:107 and Asp A:22, sulfur–π interactions with Phe A:80 and Phe A:141, and π–π stacking interactions with Tyr A:20.

Flutemetamol (2b) establishes hydrogen bonds with Thr A:227 and Lys A:48, sulfur-π interactions with Met A:256, Met A:232, and Phe A:229, and carbon–hydrogen interactions with Trp A:53, Ser A:228, Gly A:259, and Phe A:255. Flutafuranol (3b) forms a hydrogen bond with His A:107 and is further stabilized by hydrophobic interactions with Phe A:23, Phe A:80, Phe A:141, and Tyr A:20 (Sup. Fig. 3).

In the SULT1A1 catalytic pocket, PiB (1b) forms a hydrogen bond with His A:108, sulfur-π interactions with Tyr A:240 and Phe A:142, and engages in π–π stacking interactions with neighboring aromatic residues Phe A:142, Phe A:81, and Phe A:76. Flutemetamol (2a) forms a hydrogen bond with His A:149, sulfur-π interactions with Phe A:24 and Phe A:142, and hydrophobic interactions with Phe A:84, Tyr A:240, Phe A:76, and Phe A:81, while flutafuranol (3a) establishes key contacts with Lys A:48 and a network of hydrophobic interactions involving His A:108, Tyr A:240, and Phe A:81, Phe A:76, and Phe A:84 (Sup. Fig. 4).

In SULT1A3, ligands whose benzo-fused heterocycle is oriented inward relative to the catalytic site share a common interacting motif involving residues His B:149, Phe B:142, Phe B:81, Tyr B:76, and Asp B:86, reflecting a conserved network of π–π stacking, hydrophobic contacts, and polar interactions. PiB (1a) and flutemetamol (2a) share sulfur-π interactions with Phe B:24 and Phe B:81. Notably, flutafuranol (3a) adopts a particularly favorable configuration due to strong hydrogen bonding with Tyr B:139 and Lys B:106, complemented by halogen bonds with Glu B:146 and Asp B:86 (Sup. Fig. 5).

In SULT4A1, where only the catalytic site–oriented pose was retained, ligand stabilization is driven by a combination of hydrogen bonding and hydrophobic interactions (Sup. Fig. 6). PiB (1a) forms multiple hydrogen bonds with Lys A:109, Thr A:58, Lys A:55, and Tyr A:91, complemented by hydrophobic contacts with Pro A:94 and Val A:88. Flutemetamol (2a) establishes a hydrogen bond with Tyr A:19 and engages in a sulfur-π interaction with Met A:152, alongside additional π-alkyl and alkyl interactions involving Phe A:30, Pro A:28, Pro A:29, Lys A:18, and Phe A:15. Flutafuranol (3a) is stabilized by hydrogen bonds with Tyr A:142 and Tyr A:91, as well as a sulfur-π interaction with Met A:152 and hydrophobic contact with Pro A:28, indicating a balanced contribution of polar and apolar interactions within the SULT4A1 binding pocket.

Across the studied SULT isoforms, Aβ-PET tracers interact with a partially conserved set of residues that provide a balance of polar and nonpolar interactions. This recurring engagement of aromatic residues (Phe, Tyr, His) and charged side chains (Lys, His), together with consistent Tyr-mediated hydrogen bonds, is consistent with the presence of shared interactions that stabilize the ligands.

### Binding energy trajectories and RMSD profiles of Aβ PET tracers across brain-expressed SULT isoforms

Following equilibration, MD production runs (50–300 ns) were analyzed to evaluate ligand stability and interaction strength across the four SULT isoforms (Fig. 1). Complex stability was assessed by combining time-averaged binding energies with RMSD profiles. Although all three tracers exhibited binding energies within a comparable range, isoform-specific differences were observed in both mean values and fluctuation patterns. In SULT1E1 (Fig. 1A–B), all tracers displayed comparable mean binding energies.

**Figure 1.**
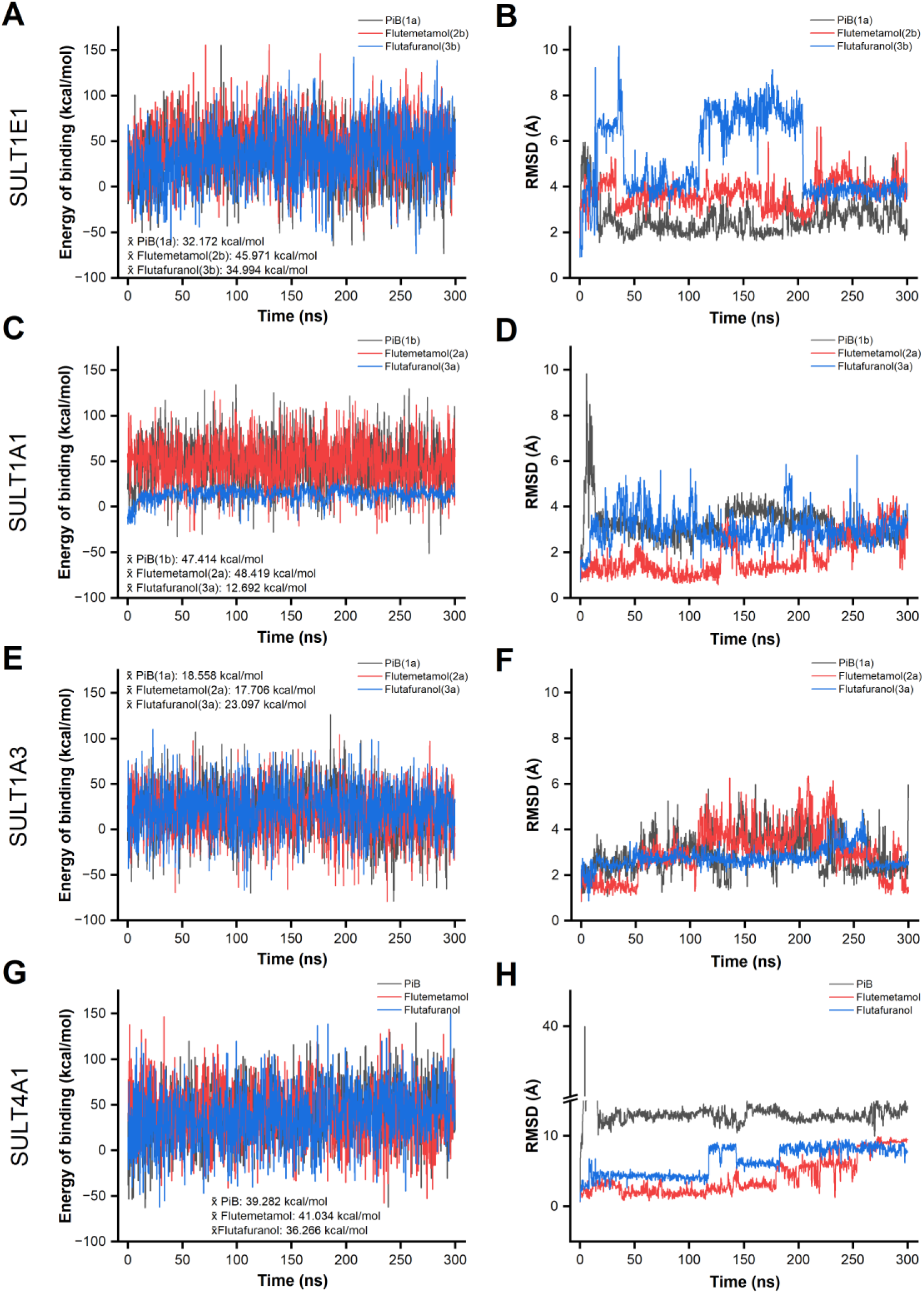
Binding energy and Root Mean Square Deviation (RMSD) profiles of Aβ PET tracers across SULT isoforms. The figure illustrates the results of 300 ns molecular dynamics (MD) simulations for PiB (black), flutemetamol (red), and flutafuranol (blue) in complex with SULT1E1 (A, B), SULT1A1 (C, D), SULT1A3 (E, F), and SULT4A1 (G, H). Panels A, C, E, G depict binding energy trajectories expressed in kcal/mol over the 300 ns simulation time. Average binding energy values for each tracer-protein complex are displayed as indicated. Panels B, D, F, H depict Root Mean Square Deviation (RMSD) profiles, expressed in Ångström (Å), calculated for the heavy atoms of the ligands relative to their initial docking positions. The Y-axis scale is adapted in panel H to accommodate the range of values observed for SULT4A1. Note that the stable accommodation of PiB within the SULT1E1 binding pocket recapitulates the experimentally established PiB-SULT1E1 interaction [23], supporting the predictive validity of the docking and MD pipeline used here and, by extension, lending confidence to the binding modes predicted for the brain-expressed isoforms SULT1A1, SULT1A3, and SULT4A1.

Nevertheless, RMSD analysis revealed distinct stability profiles: flutemetamol and PiB maintained relatively stable conformations (∼3–5 Å), whereas flutafuranol underwent a pronounced destabilization after ∼100 ns. This indicates that, despite similar mean binding energies, flutafuranol experiences significant conformational rearrangements, mirrored by the broader dispersion of its energy profile.

In SULT1A1 (Fig. 1C–D), PiB displayed low RMSD values (1–5 Å) with minimal fluctuations and relatively high mean binding energy, suggesting a stable complex. Flutemetamol showed comparable stability and binding energy, with RMSD ranging between 1 and 4 Å. However, flutafuranol, exhibited a slightly lower mean binding energy and larger RMSD (2–6 Å) with noticeable fluctuations, suggestive of a weaker and less stable interaction.

In SULT1A3 (Fig. 1E–F), PiB RMSD showed significant fluctuations (∼1.5–6 Å) despite overlapping mean binding energies with the other tracers. Flutemetamol showed a similar trend, with several fluctuations in RMSD over time. In contrast, flutafuranol maintained reduced fluctuation amplitude, suggesting a relatively stable binding mode.

In SULT4A1 (Fig. 1G–H), all tracers displayed comparable mean binding energies. However, PiB exhibited markedly high RMSD values (∼13–15 Å), indicating significant deviation from the initial binding pose. Flutemetamol showed improved stability, with lower RMSD (∼5–9 Å). Flutafuranol displayed comparable stability to flutemetamol, with similar RMSD values, supporting stable accommodation within the SULT4A1 binding pocket.

Overall, these results indicate that while the three tracers exhibit comparable mean binding energies across the SULT isoforms, their dynamic behaviors vary in an isoform-dependent manner. PiB and flutemetamol generally form stable ligand–protein complexes in SULT1E1 and SULT1A1, as reflected by relatively low RMSD values and modest fluctuations, whereas flutafuranol shows greater conformational variability in these two isoforms. This pattern shifts in SULT1A3, where PiB and flutemetamol fluctuate and flutafuranol adopts the most stable binding mode, and in SULT4A1, where PiB becomes markedly unstable while flutemetamol and flutafuranol remain comparatively stable.

## Discussion

In this study, we integrated database-guided expression analyses with molecular docking and MD simulations to assess whether brain-enriched SULTs could act as off-target binding partners for clinically relevant Aβ PET tracers. Our results extend prior observations that SULTs can interact with ThT-based compounds and support concerns about the specificity of PiB and its fluorinated derivatives in AD research and diagnostics.

Analysis of transcriptomic data obtained from highly curated, publicly available datasets identified SULT1A1, SULT1A3, and SULT4A1 as the primary SULT isoforms expressed in the human brain. Although Aβ PET tracers are primarily designed to target extracellular fibrillar amyloid deposits, their lipophilic properties and ability to diffuse across biological membranes support the possibility of transient intracellular partitioning, potentially enabling interactions with cytosolic proteins such as SULTs. Importantly, large-scale transcriptomic and proteomic datasets consistently indicate negligible baseline expression of SULT1E1 in the adult human cortex. This finding contrasts previous *in vitro* and *in vivo* observations showing that SULT1E1 is abundantly expressed in the brain [23]. This discrepancy can be explained twofold. First, by the conserved structure shared across the SULT1 isoforms [32]. Given the substantial sequence homology and overlapping substrate recognition among SULT1 isoforms, the antibody-based detection approaches employed in early studies may not fully discriminate between closely related family members, raising the possibility that previously reported SULT1E1 immunoreactivity partially reflects cross-recognition of SULT1 enzymes. Second, recent single-nucleus RNA sequencing analyses of autosomal dominant AD showed disease-associated upregulation of SULT1 in cortical cell populations [33]. This observation supports the idea that SULT1 family members can be dynamically regulated in AD brain tissue. This pattern indicates that individual SULT1 isoforms may be differentially represented depending on the pathological context, such as pro-inflammatory states [25], suggesting that tracer retention may also arise from the context-dependent expression of specific SULTs.

In this study, our *in silico* analysis confirms that ThT-based tracers can interact with multiple SULT isoforms, although notable isoform-specific differences were observed. Flutemetamol consistently adopted the most stable binding configurations across all isoforms, as evidenced by low RMSD values and favorable binding energy profiles throughout the MD simulations (Fig. 1). In contrast, the dynamic behavior of PiB and flutafuranol was isoform-dependent. In SULT1A1 and SULT1E1, PiB exhibited stable structural convergence comparable to flutemetamol, whereas flutafuranol underwent significant conformational rearrangements in SULT1E1 simulations (Fig. 1). Conversely, in SULT4A1, PiB displayed the highest RMSD and lowest stability, while flutafuranol – designed to improve specificity toward Aβ plaques [34] – maintained enhanced stability and a more favorable binding energy profile relative to PiB. While a tracer-dependent hierarchy of interaction stability emerged in SULT1A1 and SULT1E1, where PiB and flutemetamol demonstrated greater dynamic robustness than flutafuranol, this trend was notably absent in SULT1A3 and SULT4A1, where flutafuranol ranked among the more stably accommodated tracers. This pattern largely reflects the chemical design of these amyloid tracers: benzothiazole-based compounds retain planar aromatic scaffolds that promote π-mediated interactions within most enzymatic pockets. The divergent behavior observed in SULT4A1 suggests that its specific binding site architecture may be less accommodating to the PiB scaffold, whereas the benzofuran-based tracer flutafuranol, engineered to minimize non-specific binding, exhibits a comparatively reduced structural compatibility with SULT1E1 and SULT1A1 yet is stably accommodated in SULT1A3 and SULT4A1. Notably, the present analysis was restricted to tracers sharing the ThT-related benzothiazole and benzofuran scaffolds. The clinically used trans-stilbene compounds ¹⁸F-florbetapir and ¹⁸F-florbetaben possess a chemically distinct and conformationally more flexible backbone that is unlikely to be accommodated within the SULT catalytic pocket in the same manner.

Of note, the cortical distribution of SULT1A1, SULT1A3, and SULT4A1 overlaps with regions known to exhibit Aβ accumulation. Elevated cortical expression in frontal and parietal association regions – areas prominently involved in early Aβ deposition – indicates anatomical coexistence between tracer target regions and potential intracellular interaction partners [35]. Although our results do not challenge the view that extracellular fibrillar Aβ is the primary determinant of Aβ PET tracer signals, the presence of SULTs in regions involved in Aβ imaging warrants further investigation. In particular, the cerebellum – commonly used as a reference region for Aβ PET normalization due to its relative lack of Aβ deposition – exhibits robust expression of SULT isoforms, including SULT4A1. This observation provides a rationale for investigating whether SULT–tracer interactions may contribute to tracer retention in the cerebellum under physiologically relevant settings. However, whether SULT abundance, tracer concentration, and binding-site occupancy are sufficient to produce a detectable and significant contribution to PET signal remains to be experimentally determined.

The potential physiological relevance of SULT–tracer interactions is likely to be context-dependent. Neuroinflammatory states – shown to modulate SULT expression – are characterized by profound transcriptional and metabolic reprogramming affecting neurons, astrocytes, microglia, and vascular-associated cells. Imaging studies in inflammatory neurological disorders have reported increased PiB retention in regions lacking detectable fibrillar Aβ deposition, suggesting that inflammation-associated molecular environments may influence tracer behavior [25]. In this context, altered expression levels, inducible isoform shifts, or changes in intracellular accessibility of SULT enzymes could potentially modify tracer–protein interactions. However, whether such changes are sufficient to affect tracer retention *in vivo* remains unknown. Our simulations provide a structural rationale for testing this hypothesis, but do not establish a quantitative contribution of SULTs to PET signal independently of Aβ aggregation.

It is essential to acknowledge the limitations of the proposed approach. MD simulations evaluate interaction persistence within defined physicochemical parameters and do not incorporate the full complexity of *in vivo* pharmacokinetics. PET signal generation depends on tracer delivery, blood–brain barrier transport, intracellular uptake, metabolic turnover, concentration gradients, and competition with endogenous substrates. Therefore, the present findings should be interpreted as establishing structural plausibility rather than quantitative predictions of imaging signal magnitude.

Overall, the integration of multi-omics expression profiling with standardized structure-based simulations supports a model in which Aβ PET tracers exhibit conserved compatibility with multiple brain-expressed SULTs. These interactions appear to be non-catalytic and transient, reflecting general physicochemical accommodation within conserved enzyme architectures. Overall, under physiological conditions, constitutively expressed isoforms such as SULT1A1, SULT1A3, and SULT4A1 may represent potential intracellular interactors, whereas pathological states characterized by inflammatory modulation may further alter the enzymatic milieu. By linking enzyme expression patterns and dynamic structural behavior, this study provides a computational paradigm for generating experimentally testable hypotheses regarding potential intracellular interactions of Aβ PET tracers beyond their established binding to fibrillar Aβ. Determining whether these predicted interactions occur at physiologically relevant concentrations and contribute measurably to PET signal will require direct experimental and *in vivo* validation.

## Limitations

This study has several limitations that should be acknowledged. First, the present findings are based exclusively on *in silico* approaches, including molecular docking and MD simulations, which evaluate structural compatibility and interaction stability under predefined physicochemical conditions. These methods do not account for the full complexity of *in vivo* tracer behavior, including pharmacokinetics, blood–brain barrier transport, intracellular uptake, metabolic turnover, and competition with endogenous substrates. Second, while the physicochemical properties of amyloid PET tracers support their ability to cross cellular membranes, the extent to which they achieve sufficient intracellular concentrations to interact with enzymes such as SULTs remains uncertain. While direct experimental validation of the interactions with SULT1A1, SULT1A3, and SULT4A1 remains to be performed, the present pipeline correctly recovers the established PiB-SULT1E1 interaction [23, 25], providing an internal benchmark for the reported simulations. Future studies integrating experimental validation, like *in vitro* binding assays, autoradiography, or cellular uptake models, will be essential to determine the physiological relevance of the computational interactions described in our work. Third, binding free energy estimates derived from MM/PBSA calculations provide relative comparisons but do not represent accurate quantitative measures of binding affinity. Similarly, the stability observed in MD simulations does not necessarily translate into biologically relevant interactions under physiological conditions. Importantly, transcriptomic expression levels cannot be directly translated into intracellular protein abundance, tracer concentration, binding-site occupancy, or measurable PET signal. Consequently, the present study cannot determine whether SULT–tracer interactions are sufficient to influence reference region activity, SUVR normalization, or Centiloid estimation. These questions will require direct experimental validation and, ultimately, *in vivo* imaging studies. Finally, SULT4A1 is considered an atypical member of the sulfotransferase family, and its catalytic activity and endogenous substrates are not fully characterized. As such, the functional implications of tracer interactions with this isoform should be interpreted with caution.

## Author contributions

AG conceived and designed the study, supervised the experiments and wrote the manuscript. LM and RF performed database interrogation, *in silico* simulations, and drafted the manuscript. SDP and FT analyzed transcriptomic cortical mapping data. GF and SLS provided conceptual inputs and edited the manuscript. All authors approved the final version of the manuscript.

## Funding

We acknowledge financial support under the National Recovery and Resilience Plan (NRRP), Mission 6, Component 2, funded by the European Union – NextGenerationEU – PNRR-MAD-2022-12375706 - CUP: D73C22002140007

## AI statement

Large language model–based assistance (Claude Sonnet 4.6 and Opus 4.8, Anthropic) was used exclusively for language polishing and formatting. The authors are fully responsible for the scientific content, interpretation, and conclusions.

## Data availability

Processed data, including energy of binding calculations and RMSD of MD simulations for each SULT-tracer interaction are available through the Zenodo repository (https://doi.org/10.5281/zenodo.22304099) [36].

---

**Supplementary figure 1.**
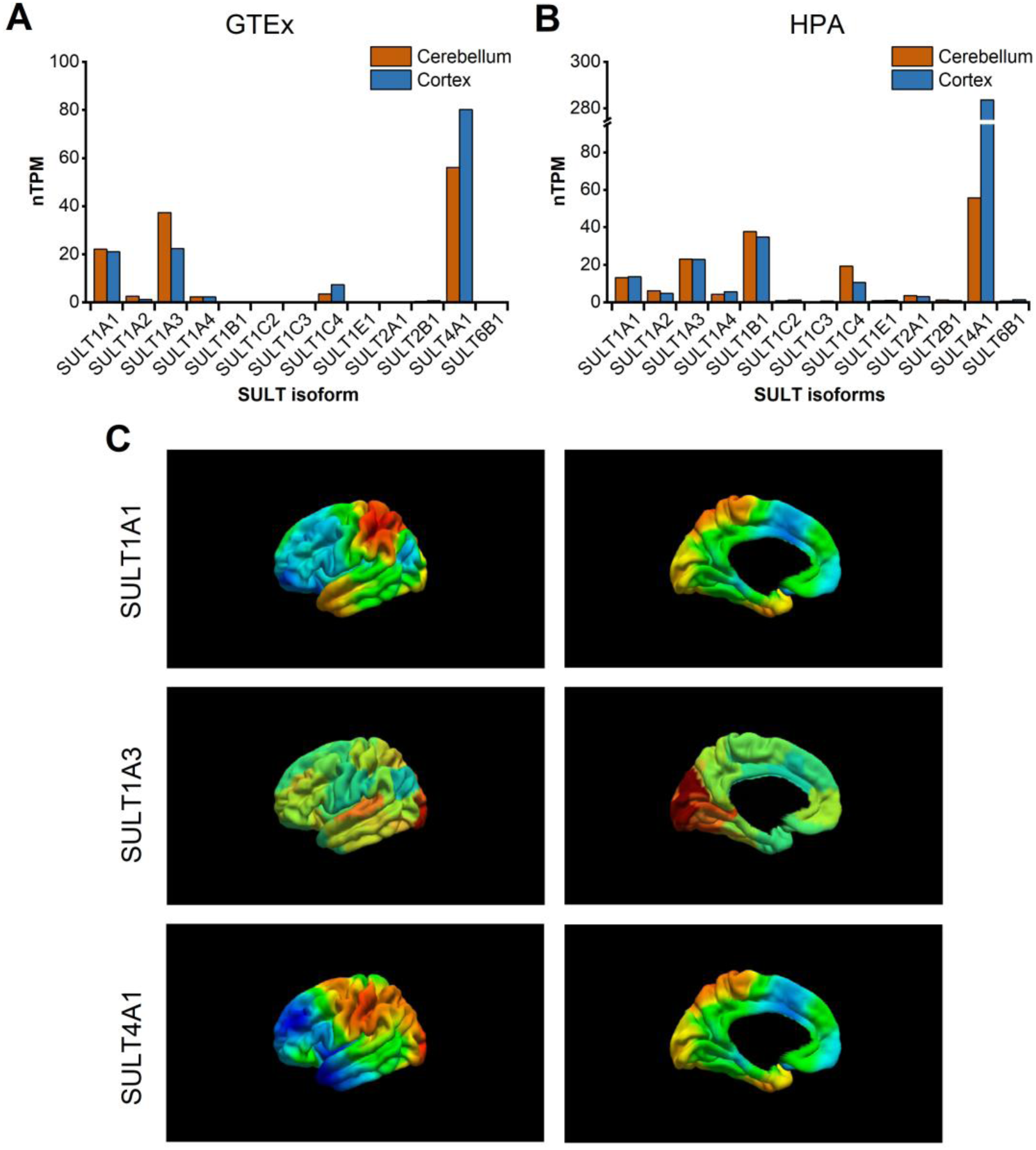
Transcriptomic expression and regional distribution of SULT isoforms in the human brain. SULT expression profiles were interrogated across two independent databases. Panels A, B show normalized expression levels (nTPM) of SULT isoforms in the cerebellum (black) and cerebral cortex (red) according to the Genotype-Tissue Expression project (GTEx; A) and the Human Protein Atlas (HPA; B). Panel C shows regional mRNA distribution maps of SULT4A1, SULT1A3, and SULT1A1 across the human brain (lateral and medial views). Color scales represent relative expression levels, with warmer colors indicating higher transcript abundance. Maps represent within-gene z-scored expression and therefore reflect relative spatial distribution rather than absolute transcript abundance across genes. Data in C were obtained from [28].

**Supplementary figure 2.**
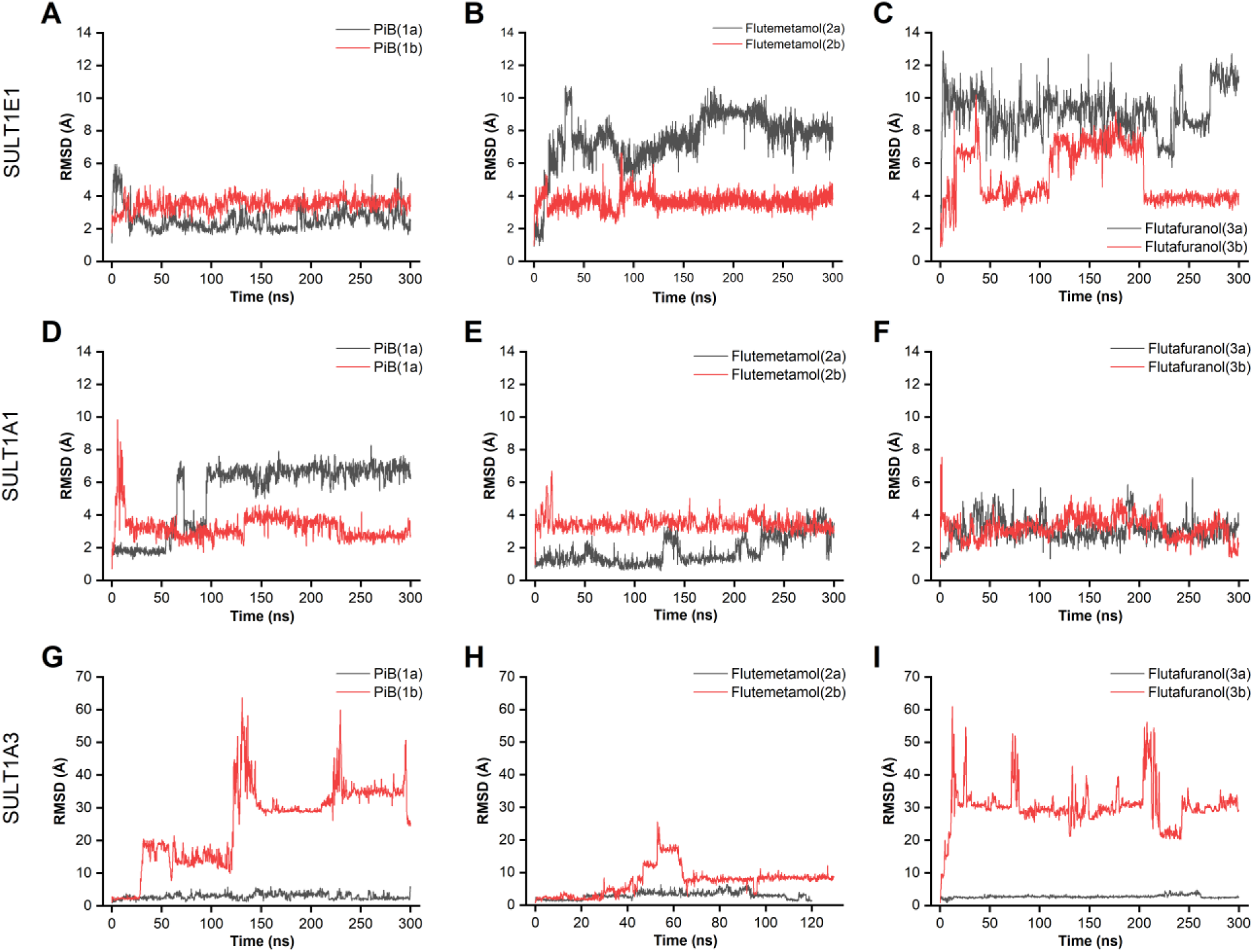
Root Mean Square Deviation (RMSD) profiles for alternative docking poses of PiB, flutemetamol, and flutafuranol. The figure displays the 300 ns molecular dynamics trajectories for two distinct docking poses (black and red lines) of each tracer across SULT1E1 (A–C), SULT1A1 (D–F), and SULT1A3 (G–I). Panels A, D, G depict RMSD profiles (Å) comparing PiB docking poses 1a (black) and 1b (red). Panels B, E, H depict RMSD profiles (Å) comparing flutemetamol docking poses 2a (black) and 2b (red). Panels C, F, I depict RMSD profiles (Å) comparing flutafuranol docking poses 3a (black) and 3b (red). The RMSD values were calculated for the heavy atoms of the ligands relative to their respective initial docking positions.

**Supplementary figure 3.**
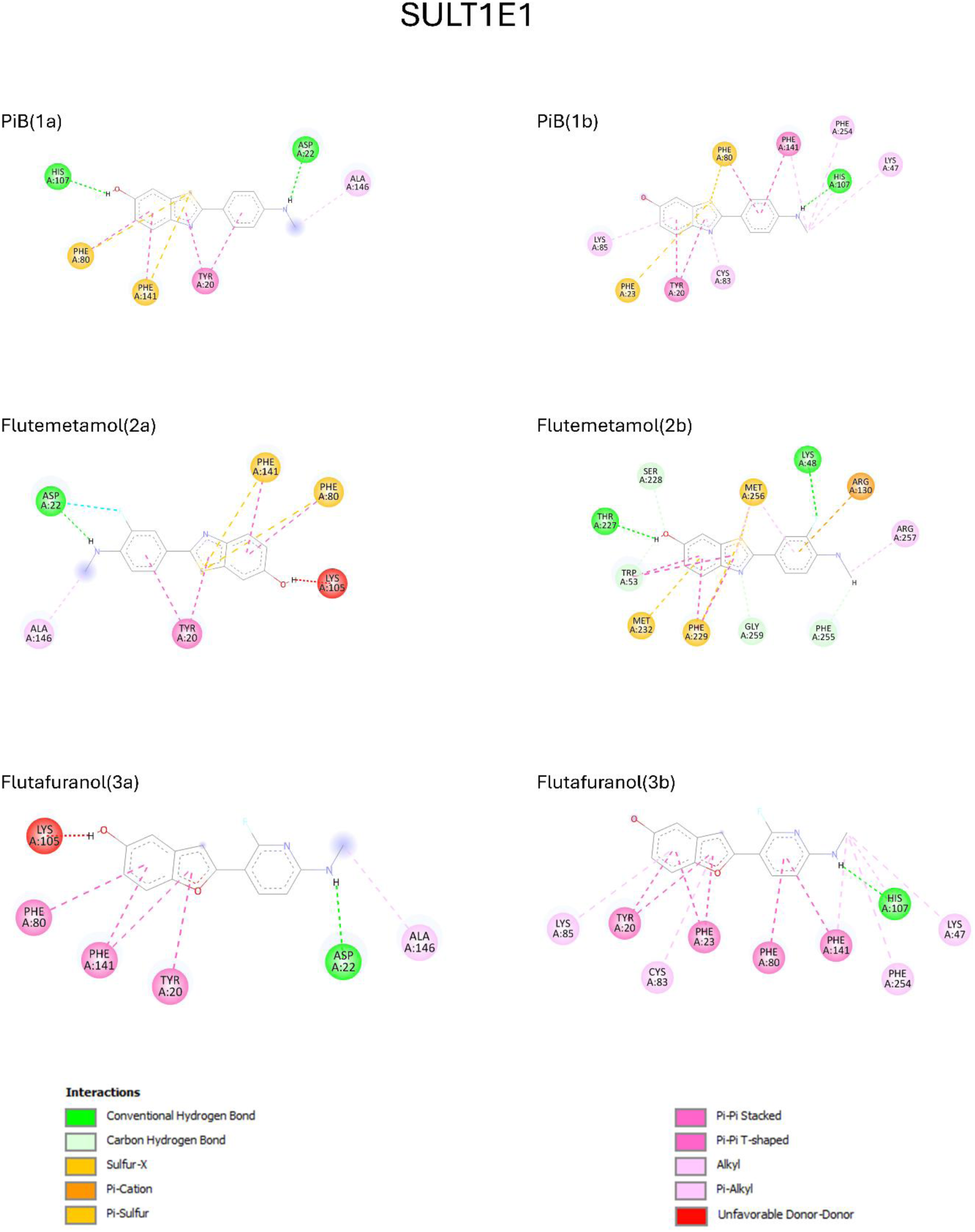
Two-dimensional interaction diagrams of alternative docking poses for SULT1E1. The figure illustrates the predicted non-covalent binding networks for PiB (1a, 1b), flutemetamol (2a, 2b), and flutafuranol (3a, 3b) within the SULT1E1 catalytic pocket. Amino acid residues are represented as labeled colored circles, with specific interactions (e.g., hydrogen bonds, π-stacking, hydrophobic contacts) indicated by color-coded dashed lines as indicated. These diagrams provide a qualitative assessment of the binding environment used to prioritize the most promising poses for subsequent 300 ns MD simulations

**Supplementary figure 4.**
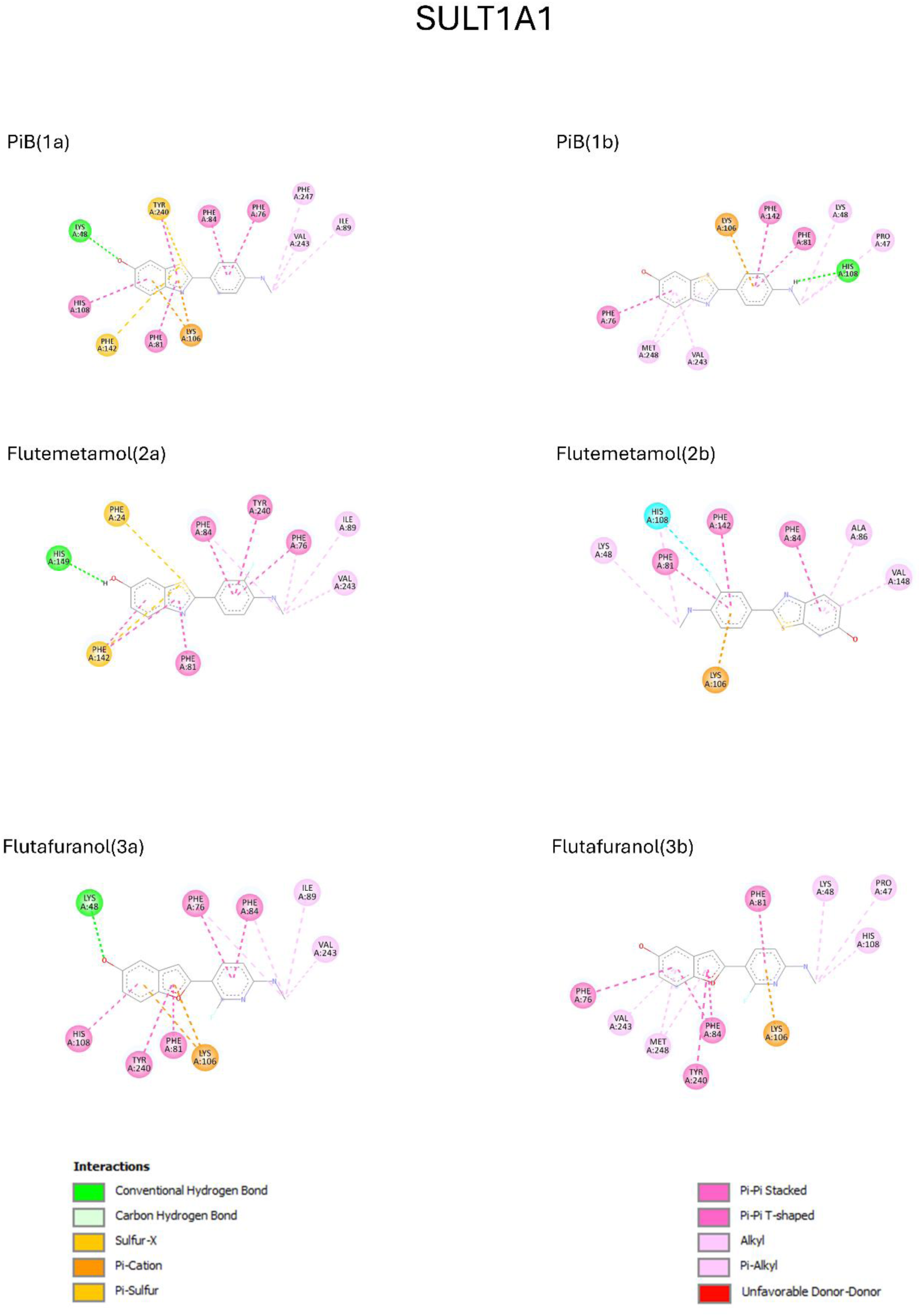
Two-dimensional interaction diagrams of alternative docking poses for SULT1A1. The figure illustrates the predicted non-covalent binding networks for PiB (1a, 1b), flutemetamol (2a, 2b), and flutafuranol (3a, 3b) within the catalytic pocket of SULT1A1. Amino acid residues are represented as labeled colored circles, with specific interactions (e.g., hydrogen bonds, π-stacking, and hydrophobic contacts) indicated by color-coded dashed lines as indicated. These diagrams provide a qualitative assessment of the binding environment used to prioritize the most promising poses for subsequent 300 ns MD simulations.

**Supplementary figure 5.**
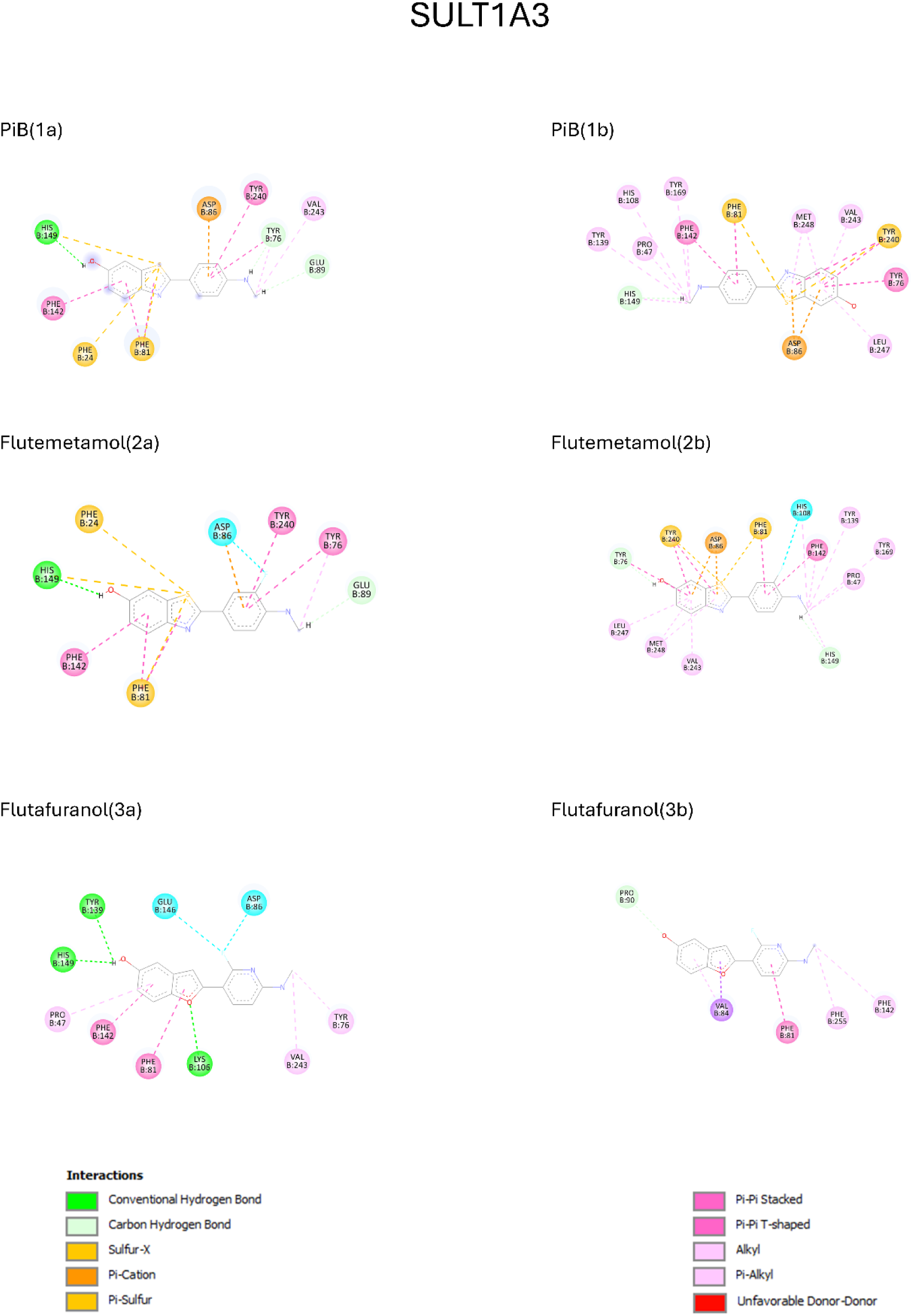
Two-dimensional interaction diagrams of alternative docking poses for SULT1A3. The figure illustrates the predicted non-covalent binding networks for PiB (1a, 1b), flutemetamol (2a, 2b), and flutafuranol (3a, 3b) within the catalytic pocket of SULT1A3. Amino acid residues are represented as labeled colored circles, with specific interactions (e.g., hydrogen bonds, π-stacking, and hydrophobic contacts) indicated by color-coded dashed lines as indicated. These diagrams provide a qualitative assessment of the binding environment used to prioritize the most promising poses for subsequent 300 ns MD simulations.

**Supplementary figure 6.**
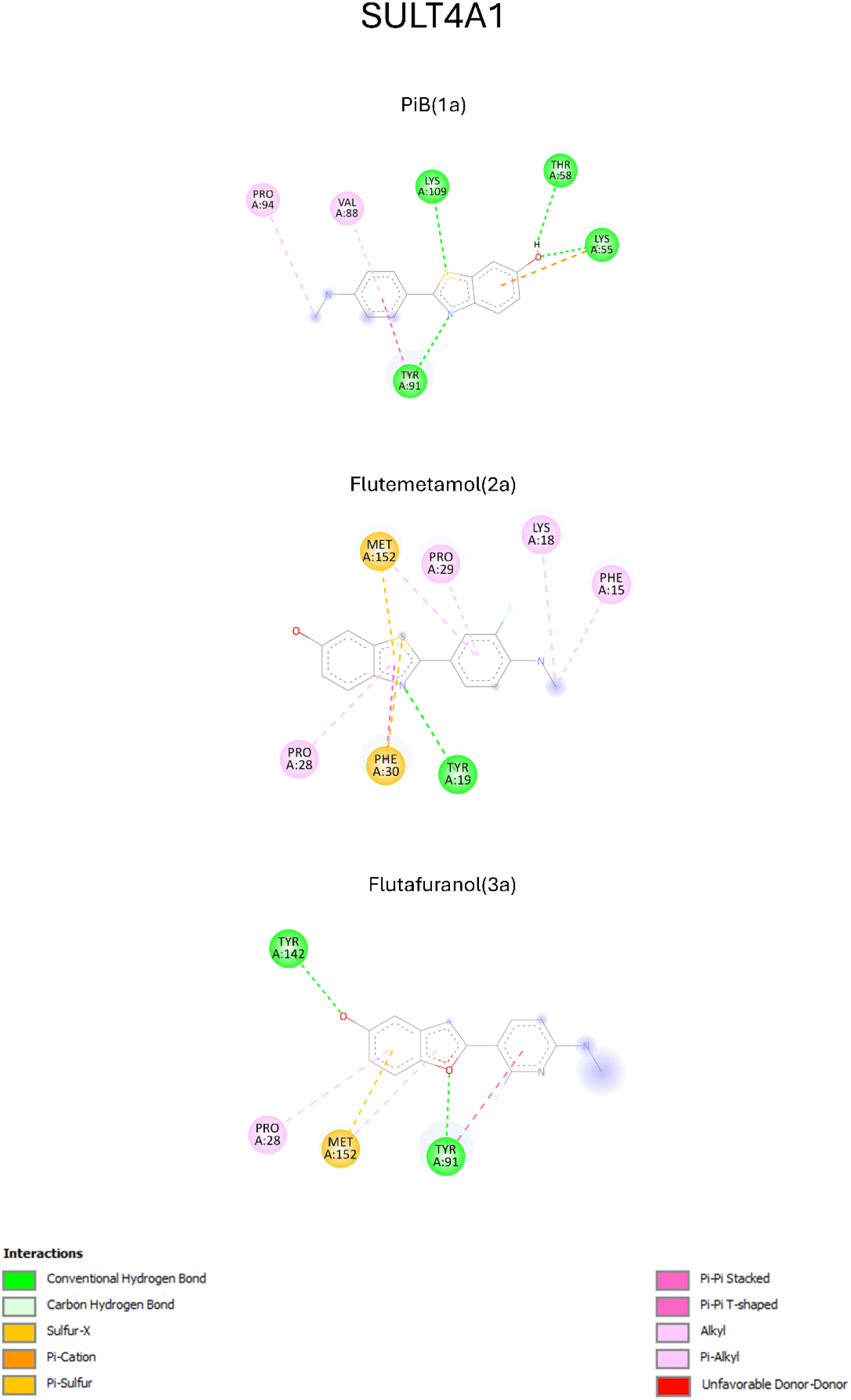
Two-dimensional interaction diagrams of docking poses for SULT4A1. The figure illustrates the predicted non-covalent binding networks for PiB (1a), flutemetamol (2a), and flutafuranol (3a) within the catalytic pocket of SULT4A1. For this isoform, a single docking pose was identified for each tracer and further investigated through 300 ns molecular dynamics simulations. Amino acid residues are represented as labeled colored circles, with specific interactions (e.g., hydrogen bonds, π-stacking, and hydrophobic contacts) indicated by color-coded dashed lines as indicated. These diagrams provide a qualitative assessment of the initial binding environment for each ligand-protein complex.

**Supplementary table 1.** Statistical analysis of RMSD trajectories for alternative docking poses. The table summarizes the RMSD parameters calculated over 300 ns molecular dynamics simulations for PiB,flutemetamol, and flutafuranol in complex with the SULT1E1, SULT1A1, and SULT1A3 isoforms. For each ligand-protein complex, two alternative docking poses (indicated as 1a/1b, 2a/2b, or 3a/3b) are compared. The reported values include the mean of RMSD (Mean, Å), the standard deviation (SD, Å), and the slope of the linear regression (Slope) derived from the RMSD vs. time plots. These descriptors were employed to evaluate the structural convergence and stability of each binding mode.

|  | Average | SD | Slope |
| --- | --- | --- | --- |
| SULT1E1_PiB(1a) | 2.4992 | 0.5621 | 0.0030 |
| SULT1E1_PiB (1b) | 3.5836 | 0.3756 | 0.0008 |
| SULT1E1_Flutemetamol(2a) | 8.33499 | 0.70680 | -0.01455 |
| SULT1E1_Flutemetamol(2b) | 3.69058 | 0.35198 | 0.00158 |
| SULT1E1_Flutafulanol(3a) | 9.0512 | 1.2939 | 0.0045 |
| SULT1E1_Flutafulanol(3b) | 5.1709 | 1.6252 | -0.0056 |
| sult1A1_PiB(1a) | 6.0873 | 1.3701 | 0.0119 |
| sult1A1_PiB(1b) | 3.1421 | 0.5494 | -0.0007 |
| SULT1A1_Flutemetamol(2a) | 1.8976 | 0.8907 | 0.0088 |
| SULT1A1_Flutemetamol(2b) | 3.4642 | 0.3650 | -0.0005 |
| sult1A1_Flutafulanol(3a) | 3.0812 | 0.6364 | -0.0012 |
| sult1A1_Flutafulanol(3b) | 3.3035 | 0.6556 | -0.0021 |
| sult1A3_PiB(1a) | 3.0528 | 0.8707 | -0.0019 |
| sult1A3_PiB(1b) | 28.2711 | 9.8976 | 0.0928 |
| SULT1A3_Flutemetamol(1a) | 3.2997 | 0.9578 | -0.0010 |
| SULT1A3_Flutemetamol(1b) | 8.8197 | 3.5457 | -0.0028 |
| sult1A3_Flutafulanol(1a) | 2.8196 | 0.4037 | 0.0007 |
| sult1A3_Flutafulanol(1b) | 30.3167 | 5.3244 | -0.0042 |

## Notes

### Competing Interest Statement

The authors have declared no competing interest.

### Summary of Updates

Softned some conclusions; improved figure quality and resolution.

