## Supplementary figures and images for "Analysis of the off-target interaction of amyloid PET tracers with human brain sulfotransferases"

### Supplementary Figure 1

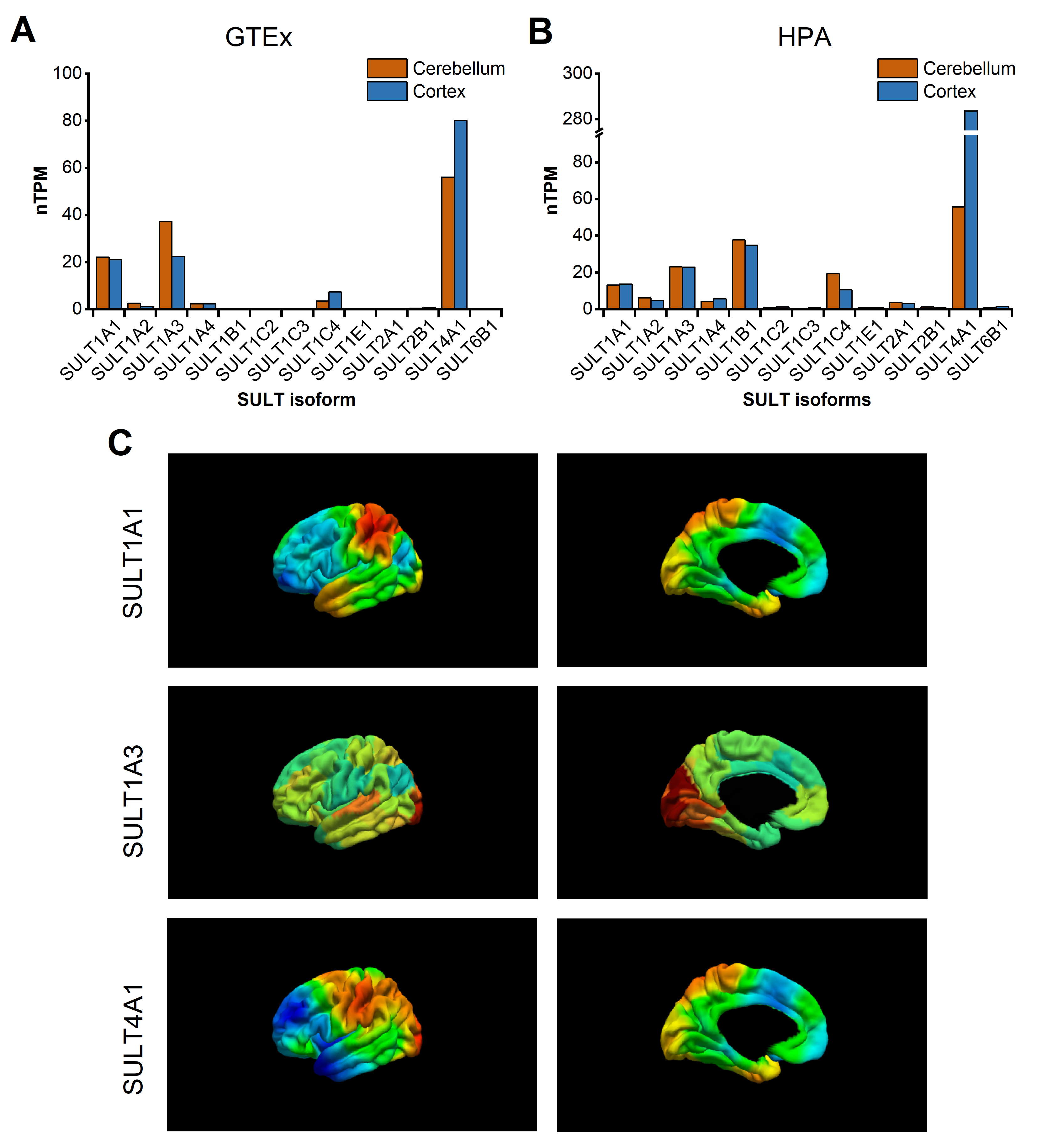

### Supplementary Figure 2

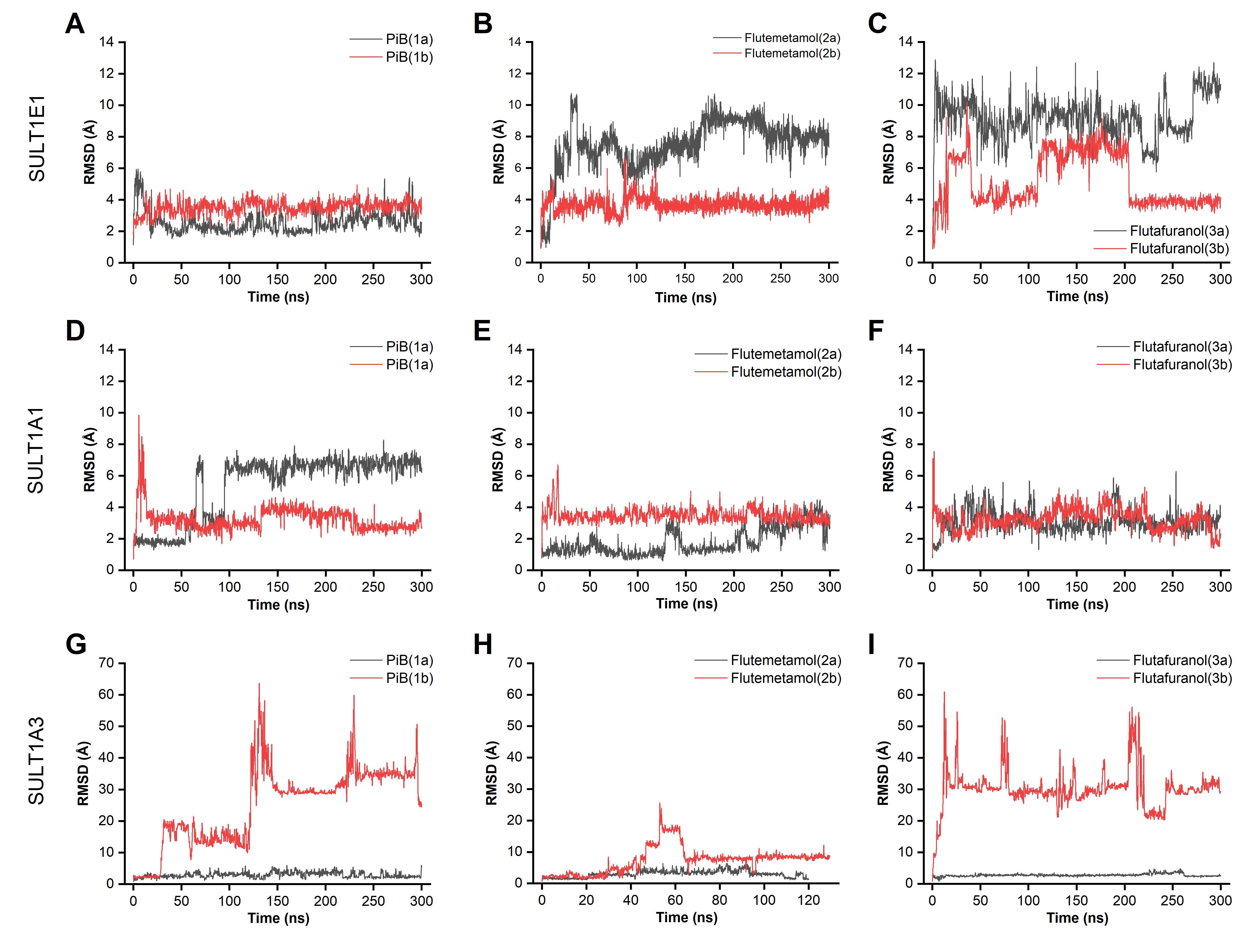

### Supplementary Figure 3

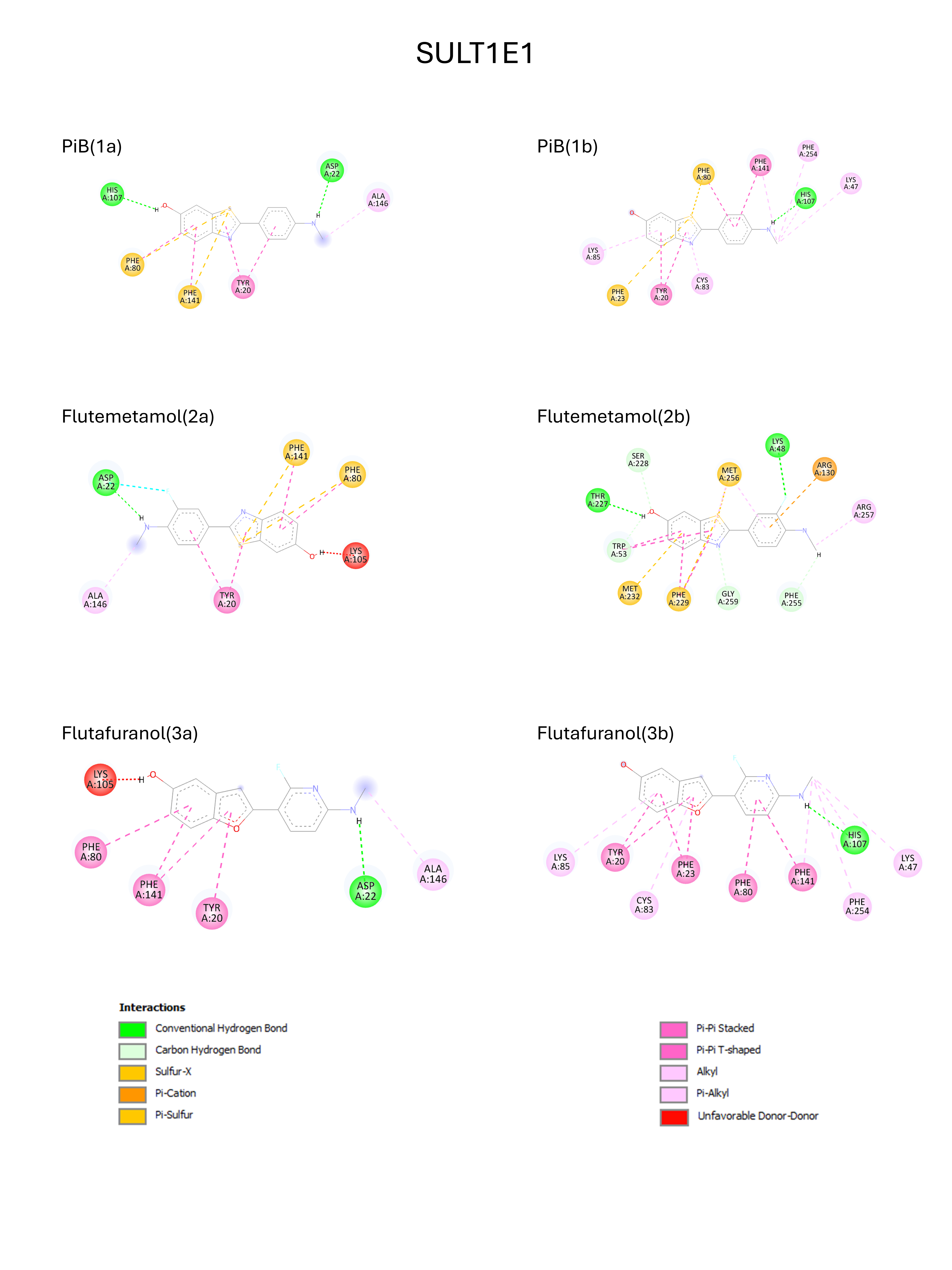

### Supplementary Figure 4

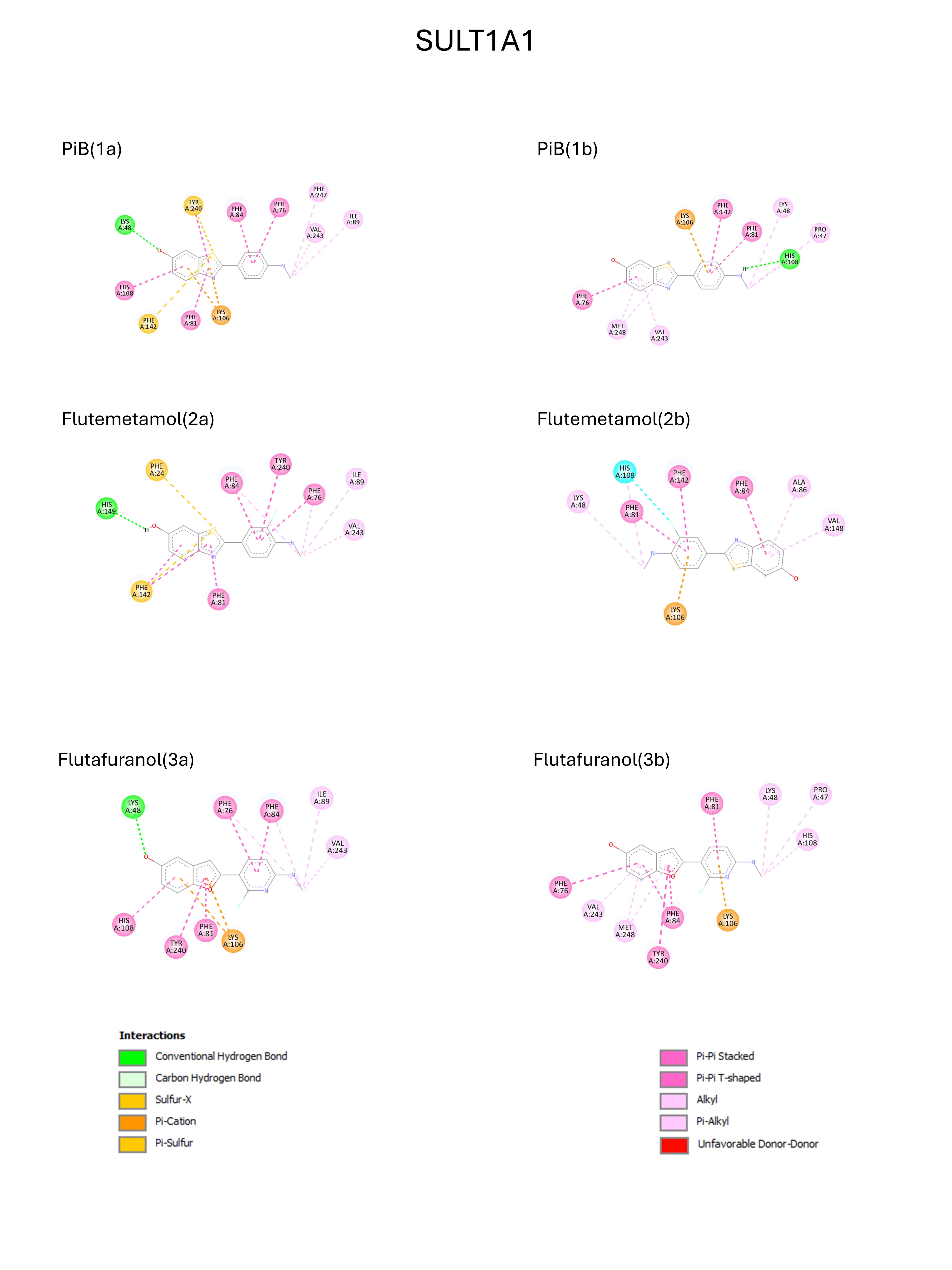

### Supplementary Figure 5

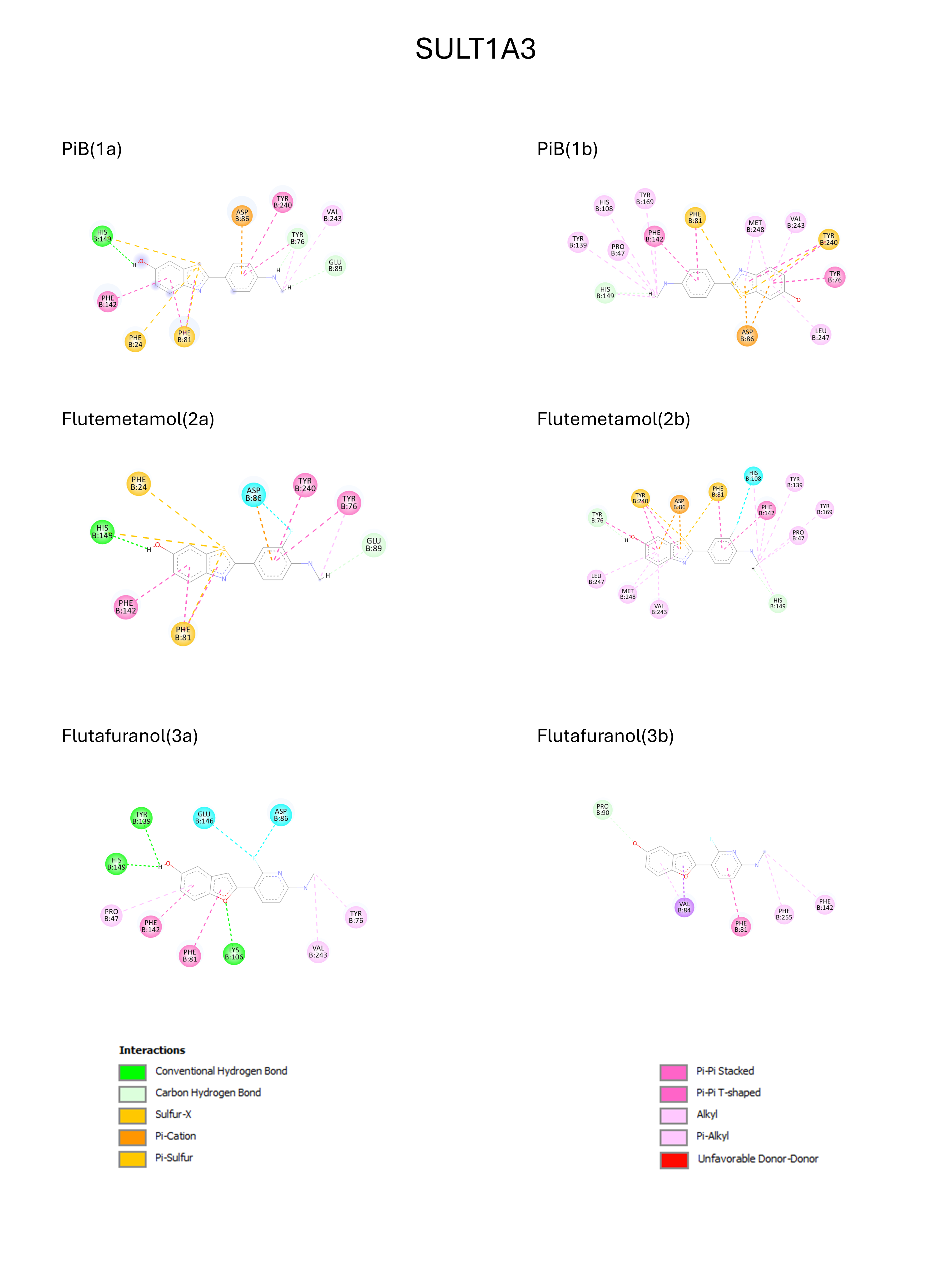

### Supplementary Figure 6

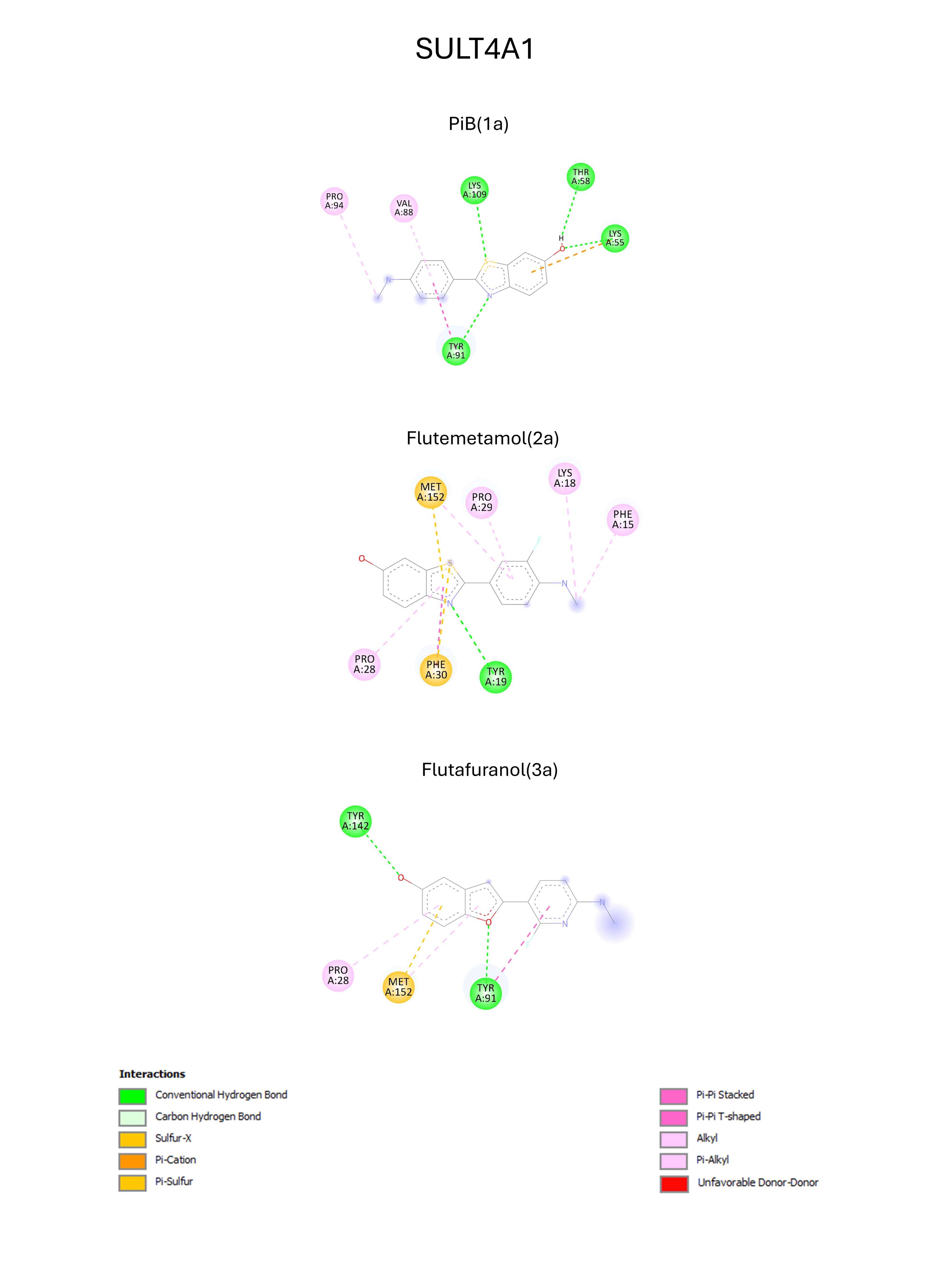

### Supplementary Table 1

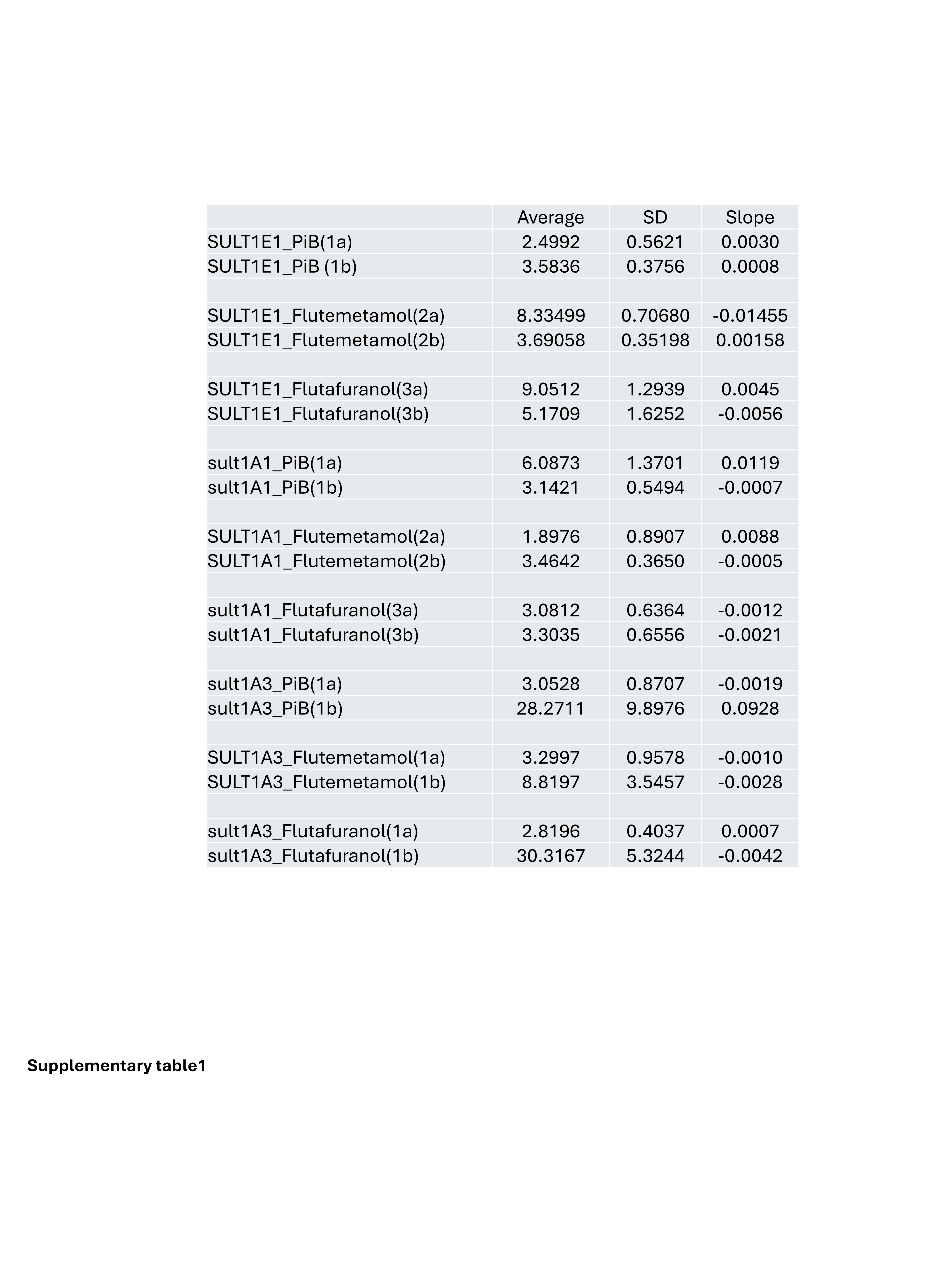
